# Whole genome sequencing reveals the emergence of a *Pseudomonas aeruginosa* shared strain sub-lineage among patients treated within a single cystic fibrosis centre

**DOI:** 10.1101/261586

**Authors:** Bryan A. Wee, Anna S. Tai, Laura J. Sherrard, Nouri L. Ben Zakour, Kirt R. Hanks, Timothy J. Kidd, Kay A. Ramsay, Iain Lamont, David M. Whiley, Scott C. Bell, Scott A. Beatson

**Author notes:** These authors contributed equally to this work (Bryan A. Wee, Anna S. Tai). Email addresses Bryan A. Wee, Anna S. Tai, Laura J. Sherrard, Nouri L. Ben Zakour, Kirt R. Hanks, Timothy J. Kidd, Kay A. Ramsay, Iain Lamont, David M. Whiley, Scott C. Bell, Scott A. Beatson.

## Abstract

**Background:** Chronic lung infections by *Pseudomonas aeruginosa* are a significant cause of morbidity and mortality in people with cystic fibrosis (CF). Shared *P. aeruginosa* strains, that can be transmitted between patients, are of concern and in Australia the AUST-02 shared strain is predominant in individuals attending CF centres in Queensland and Western Australia. M3L7 is a multidrug resistant sub-type of AUST-02 that was recently identified in a Queensland CF centre and was shown to be associated with poorer clinical outcomes. The main aim of this study was to resolve the relationship of the emergent M3L7 sub-type within the AUST-02 group of strains using whole genome sequencing.

**Results:** A whole-genome core phylogeny of 63 isolates indicated that M3L7 is a monophyletic sub-lineage within the context of the broader AUST-02 group. Relatively short branch lengths connected all of the M3L7 isolates. A phylogeny based on nucleotide polymorphisms present across the genome showed that the chronological estimation of the most recent common ancestor was around 2001 (± 3 years). SNP differences between sequential M3L7 isolates collected 3-4 years apart from five patients suggested both continuous infection of the same strain and cross-infection of some M3L7 variants between patients. The majority of polymorphisms that were characteristic of M3L7 (i.e. acquired after divergence from all other AUST-02 isolates sequenced) were found to produce non-synonymous mutations in virulence and antibiotic resistance genes.

**Conclusions:** M3L7 has recently diverged from a common ancestor indicating descent from a single carrier at a CF treatment centre in Australia. Both adaptation to the lung and transmission of M3L7 between adults attending this centre may have contributed to its rapid dissemination. The study emphasises the importance of clinical management in controlling the emergence of shared strains in CF.

## Background

Cystic fibrosis (CF) is the most common recessively lethal inherited disease in people of European ancestry. The majority of mortality and morbidity in people with CF is caused by chronic lung infection with *Pseudomonas aeruginosa* [1]. During chronic infection *P. aeruginosa* adapts to the CF airway microenvironment, which promotes multiple phenotypic and genotypic changes including enhanced resistance to antibiotics, excessive exopolysaccharide production, auxotrophy, auxotrophic metabolism, hypermutability and the loss of motility [2-8]. This evolution strategy has been found to occur in parallel in different *P. aeruginosa* strains, suggesting that these pathoadaptive modifications are important in the transition from an opportunistic pathogen to a specialised pathogen of diseased human lungs [8].

Person-to-person transmission of airway-adapted *P. aeruginosa* strains has also been reported with the acquisition of some “shared strains” correlated with adverse clinical outcomes [9-13]. AUST-02 is a prevalent shared strain in CF centres around Australia and is the dominant strain in Queensland infecting approximately 40% of patients infected with *P. aeruginosa* [10, 14, 15]. A recent surveillance study also described the detection of an AUST-02 strain sub-type in approximately 5% of patients attending a single CF treatment centre in Brisbane (Queensland) [16]. This sub-type could be distinguished from all other AUST-02 strains by a unique *mexZ* (*mexZ-3,* M3) and *lasR* (*lasR*-7, L7) genotype and therefore, was designated M3L7 [16]. The M3L7 sub-type is of particular importance given that, compared to other AUST-02 and *P. aeruginosa* strains, it is associated with enhanced antibiotic resistance and poorer clinical outcomes (e.g. higher hospitalisation requirements and increased risk of lung transplantation or death) [16].

The aim of this study was to use whole genome sequencing (WGS) to reconstruct the population structure and resolve the relationship of the M3L7 sub-type within the AUST-02 group of strains. The analyses revealed that the M3L7 sub-type has recently diverged from a common ancestor suggesting descent from a single founder within a rapidly growing CF centre population. Genetic mutations exclusive to the M3L7 sub-type were identified and may have aided adaptation to the CF airway microenvironment prior to its dissemination.

## Methods

### Bacterial isolates and whole genome sequencing

*P. aeruginosa* isolates (n=25) encompassing the majority of M3L7 isolates (named AUS934 to AUS958) that were recently described, and one isolate considered an outgroup (AUS970) were chosen for sequencing (Additional File 1: Table S1) [16]. These isolates originated from expectorated sputum (collected in 2007, 2008 and 2011) provided by adults with CF (n=20), who attended The Prince Charles Hospital Adult CF centre in Brisbane for their clinical care [16]. Preparation of genomic DNA for WGS was undertaken using the UltraClean^®^ Microbial DNA Isolation Kit as described previously [17]. Library preparation (Truseq), qPCR (TapeStation, Agilent Genomics) and WGS using the Illumina HiSeq 2500 platform with 100 bp paired-end read chemistry were carried out by the Australian Genome Research Facility, Melbourne, Australia.

In order to reconstruct the M3L7 population structure, a further 37 AUST-02 genomes that were previously sequenced (as part of an ongoing AUST-02 population genetic diversity study) were included in the analyses (Additional File 1: Table S1).

### Genome mapping and assembly

Reads were taxonomically assigned with Kraken (v0.10.4) to check for contamination and trimmed using Nesoni clip (v0.128) to filter out adapter sequences and low-quality regions [18, 19]. Reads were mapped to the *P. aeruginosa* PAO1 reference genome (NC_002516) using Shrimp as implemented in Nesoni (v0.128) [18, 20, 21]. The PAO1 genome was chosen as a reference due to its high quality, expert-curated annotation [21]. SNPs and small insertions or deletions (indels) shorter than the read length were called using Nesoni.

Genomes were assembled using Velvet (v1.2.10) and VelvetOptimiser (v2.2.5) [22, 23]. Assembled contigs were reordered against PAO1 using Mauve (v2.4.0) and annotated with Prokka (v1.10) [24, 25]. Gene annotations from PAO1 were used as the primary reference.

### Phylogenetic analysis

An alignment of 30,811 core SNPs obtained from mapping against PAO1 was used to reconstruct the phylogeny of the 63 AUST-02 genome sequences. RAxML (v8.1.15) was used to estimate the Maximum Likelihood tree with the rapid bootstrap analysis option (`-f a`) and GTRGAMMA model of nucleotide substitution with a correction for ascertainment bias (`-m ASC_GTRGAMMA --asc-corr lewis`) [26]. A resolved phylogeny of the M3L7 genome sequences (generated in this study) was constructed from a SNP matrix of 364 SNPs with the same settings as above. Phylogenetic trees were viewed and explored using Dendroscope (v3.4.1) and FigTree (v1.4.2) [27, 28]. Minimum spanning trees were generated using the goeBURST Full MST function in Phyloviz [29].

### BEAST analysis

To determine the emergence of the M3L7 sub-type, Bayesian inference of the evolutionary rates was conducted using BEAST 1.8.2 [30]. As input a set of 183 SNPs specific for M3L7 was used, excluding two hypermutator isolates (AUS937 and AUS938) and the AUS970 outgroup. Regions of clustered SNPs, where at least three SNPs were found within 10bp of each other, were also removed. Among the different combinations of the molecular clock model (strict and constant relaxed lognormal), substitution model (HKY, GTR) and population size change (coalescent constant and exponential growth) models, the preferred combination of parameters selected based on stepping stone sampling was strict molecular clock, HKY substitution model with four discrete gamma-distributed rate categories, and exponential population size change. Markov Chain Monte Carlo generations were run in triplicate for 50 million steps, sampling every 5,000 steps, to ensure convergence and an ESS value > 200 for all parameters. Replicate runs were combined using LogCombiner with a 10% burn-in and maximum credibility trees reporting mean values were created using TreeAnnotator.

### Comparative genomic analyses

Comparative genomic analyses were performed using Parsnp, Gingr, BRIG (BLAST Ring Image Generator), Roary (v3.4.2), ACT (Artemis Comparison Tool) and BLAST [31-34]. Large genomic differences were investigated using PHAST (PHAge Search Tool) and Roary [33, 35]. The effect of amino acid substitutions (functionally important or no change) were predicted *in silico* using PROVEAN (Protein Variation Effect Analyzer) [36].

## Results and Discussion

### Isolates selected for WGS in this study

One to three *P. aeruginosa* isolates were prospectively collected from single sputum specimens obtained annually for culture as part of the Australian Clonal *P.aeruginosa* in Cystic Fibrosis (ACPinCF) study [10] and a systematic surveillance study was subsequently conducted to assess the prevalence of M3L7 between 2007 and 2011 [16]. M3L7 was identified in 28/509 (5.5%) *P. aeruginosa* isolates from 13/170 (7.6%) patients in 2007-2009 and in 21/519 (4.0%) *P. aeruginosa* isolates from 11/173 (6.4%) patients in 2011 [16]. Twenty-five of these isolates were selected from all patients (patients 24 to 42; Additional File 1: Table S1) infected with M3L7 for WGS in this study.

Of the 13 patients identified with M3L7 in 2007-2009, five underwent lung transplantation and one moved interstate by 2011. Therefore, no follow-up isolates were available. Of the seven patients who had M3L7 infection in 2007 and had samples collected in 2011, five remained infected in 2011, one no longer had the M3L7 subtype detected, while another tested positive by the M3 allele-specific PCR, but the *lasR* sequence could not be determined because of suboptimal sequence quality (this isolate, AUS947, was subsequently confirmed as M3L7 by WGS). A further five patients acquired M3L7 in 2011 and were identified as incident cases (these patients were infected with other strains in 2007). Finally, one patient infected with M3L7 in 2011 had no previous strain-typing data available.

One M3L7 isolate was randomly selected per patient at each time-point (2007-2009 and 2011) for WGS and comprised: i) 13 M3L7 isolates from 13 patients in 2007-2009; ii) six M3L7 isolates from six patients with persistent M3L7 infection in 2011; iii) five M3L7 isolates from five incident cases in 2011; and iv) one M3L7 isolate from a patient with no previous *P. aeruginosa* strain typing data collected in 2011. One further *P. aeruginosa* isolate from a patient (patient 43; Additional File 1: Table S1) in 2007 which contained the M3 *mex*Z allele and a non-L7 *lasR* genotype was included as an outgroup.

### M3L7 is a distinct sub-lineage of the AUST-02 shared strain

On the basis of the whole-genome core phylogeny, the 63 AUST-02 isolates form two major discrete lineages (clades), M2 (n=34) and M3 (n=29), consistent with their possession of *mexZ-2* (codon substitution, A38T) or *mexZ-3* (codon substitution, T12N) alleles, respectively (Figure 1 and Additional File 2: Figure S1). The M3L7 isolates (n=26; including 25 isolates sequenced here and a previously sequenced AUST-02 genome, AUS22) form a monophyletic sub-lineage of AUST-02 within the M3 clade that has diverged from all other AUST-02 isolates sequenced to date (Figure 1). Three isolates (AUS853; AUS854; AUS970) within the M3 clade form deep-branching relationships at the base of the lineage and do not harbour the *lasR-7* allele (L7: 1 bp deletion, 438delG) that defines the M3L7 sub-type. Of the available AUST-02 sequences, isolate AUS970 (sequenced in this study) represented the AUST-02 genome that was most closely related to the M3L7 sub-lineage. In addition to the *mexZ*-*3* allele (M3), AUS970 carried a wild-type *lasR-1* (L1) allele (therefore named M3L1).

**Figure 1.**
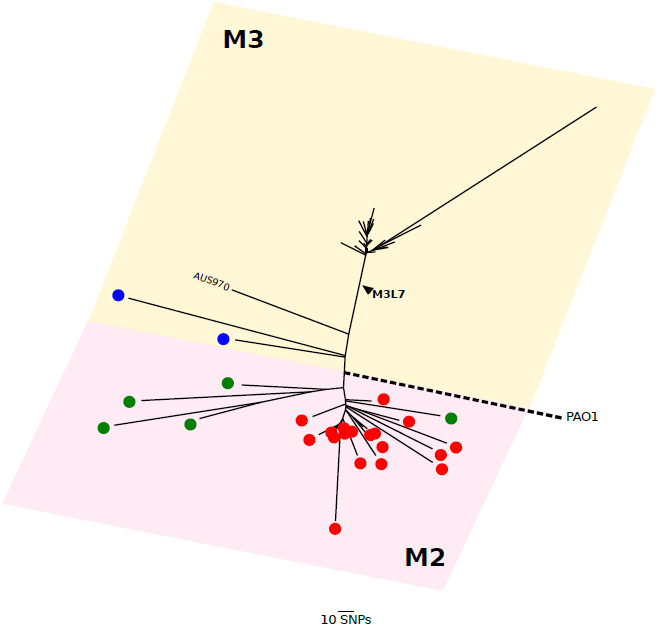
Radial phylogeny of the AUST-02 genomes. The relationship of the M3L7 sub-lineage to other sequenced AUST-02 genomes from patients attending CF centres in Brisbane/Queensland (Red), Perth/Western Australia (Green) and Sydney/New South Wales (Blue) is shown. The outgroup (AUS970) is indicated. The major clades (M2 and M3) are represented by pink and yellow shaded boxes and are defined by different *mexZ* alleles (M2, A38T; M3, T12N). The scale bar represents 10 nucleotide substitutions. Phylogeny was reconstructed estimated from an alignment of 30,811 core genome SNPs (relative to PAO1) using RAxML.

### M3L7 diverged recently from other AUST-02 shared strains

The M3L7 sub-lineage expanded recently following a long period of divergence from the M2 sub-lineage according to the relatively short branch lengths connecting the M3L7 isolates and their relative distance from the root (Figure 1). Using BEAST analysis (Figure 2), we estimated that the most recent common ancestor (MCRA) of M3L7 emerged around 2001 (± 3 years). This is approximately six years prior to the first isolation of an M3L7 sub-type (2007) in people with CF in Brisbane (Additional File 3: Figure S2) [16]. Of note, this time period also corresponds with a relatively high annual increase (approximately 10-15%) in the adult CF population at The Prince Charles Hospital (Additional File 3: Figure S2). This situation, combined with limited capacity to segregate all patients, particularly when admitted to an inpatient ward, may have contributed to shared-strain infections. Taken together, these results support a single founder scenario in which a carrier of the M3L7 sub-type acted as a donor within a CF centre with subsequent rapid dissemination in the resident CF population. Notably, both sub-lineages of AUST-02 are prevalent in Queensland, suggesting that their MCRA originated in this state; however, we cannot rule out independent introductions of M3L7 and M2 founders into the local CF population from outside the same geographical region (state).

**Figure 2.**
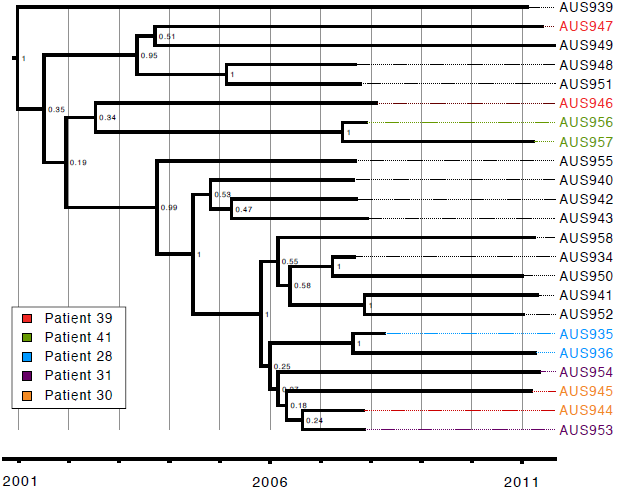
Time-calibrated phylogeny of the M3L7 sub-lineage. Ancestral reconstruction was performed using BEAST 1.8.2 based on a 183 bp non-recombinant SNPs alignment for the 23 non-hypermutator M3L7 strains (sequenced in the current study) isolated between 2007 and 2011, with HKY substitution-, strict clock-, and exponential population tree-models preferred. Posterior probability support is indicated for each node. Paired samples from a patient are coloured according to the legend depicted on the bottom left corner. The x-axis represents the years between 2001-2011.

### Resolving the within-patient relationships of M3L7 provides evidence for both continuous infection and person-to-person transmission

To investigate genomic evidence for continuous infection within patients and person-to-person transmission, we examined the pattern of SNP differences between isolates in more detail. As described previously, the dataset included six pairs of M3L7 isolates (Additional File 2: Figure S1) collected from people with CF at two time-points in 2007 or 2008 and in 2011 [16].

A pair of M3L7 isolates (AUS937 and AUS938) from patient 38 had accumulated a much higher number of SNPs compared to other M3L7 isolates, as indicated by their relatively longer branches in the phylogeny (Additional File 2: Figure S1). Further analysis revealed that these two isolates had acquired independent non-synonymous mutations within the *mutS* gene (AUS937, L341P; AUS938, 1 bp deletion [1076delC]), encoding a DNA mismatch repair protein [37], and were predicted to be deleterious to protein function based on an *in silico* analysis [36]. Mutations in *mutS* are associated with hypermutation, which frequently occurs during chronic *P. aeruginosa* infection of the CF airway [38, 39].

After removal of the AUS937 and AUS938 isolates from the analysis, the phylogeny of the M3L7 sub-lineage could be resolved further: a maximum of 47 core SNPs separated the most divergent (AUS941 and AUS947) isolates (Additional File 4: Figure S3 and Additional File 5: Figure S4). The closest sequential within-patient isolates (AUS956 and AUS957 from patient 41) differed by only two core SNPs demonstrating a remarkably high degree of genome stability over a 4-year period (Additional File 4: Figure S3). Within-patient isolates from patients 41 and 28 also grouped together with high bootstrap support on the phylogeny, which is consistent with continuous M3L7 infection across the two sampling time-points. Our analyses also revealed that a very close relationship existed between the early and late isolates from patients 30 and 31, which could also be due to continuous infection of the same strain (Additional File 4: Figure S3).

In contrast, based on the Maximum-Likelihood phylogeny, the second isolate (2011) cultured from patient 39 was grouped together with isolates from other patients (patient 36; patient 37; patient 40) instead of only the earlier 2007 isolate (Additional File 4: Figure S3). This indicates the possibility of multiple M3L7 cross-infection events as has been suggested for other shared strains [40], which may occur via the airborne route or during socialisation between patients [41, 42]. Pairwise SNP comparisons between M3L7 isolates also shows a possible transmission pathway with isolates AUS946 and AUS944 from patient 39 and patient 30, respectively, being the most likely source of cross-infections (Additional File 5: Figure S4).

AUS22 (an M3L7 isolate sequenced as part of a different study) and AUS943 were both isolated from patient 32 just six days apart. However, these two isolates were more closely related to isolates from other patients than to each other, which is also suggestive of direct or indirect cross-infection between those patients (Additional File 2: Figure S1). This finding highlights that short-term within-patient diversity of shared strains during chronic infection (e.g. [17]) needs to be considered in light of the *P. aeruginosa* lung diversity of the local CF population as a whole.

The resolution of WGS data of multiple isolates from individual patients, combined with social interaction data will enable future studies to distinguish between continuous infection or recent acquisition of M3L7 variants amongst the CF population [14].

### M3L7 is characterised by an accumulation of non-synonymous mutations in critical pathways

Clonal lineages are expected to accumulate mutations that enable adaptation of the bacterium to a specific environmental niche of a human host [8]. A total of 44 shared SNPs and nine shared indels were acquired after divergence of the M3L7 sub-lineage from all other AUST-02 isolates of the M2 and M3 clades. Thirty-five SNPs were non-synonymous (80%), resulting in a change of the amino acid sequence including two premature stop codons (Additional File 6. Table S2). Four indels produced in-frame mutations, whilst five indels caused a shift in the reading frame (Additional File 6: Table S2).

Genes containing non-synonymous SNPs and indels exclusive to the M3L7 sub-lineage (n=43) were subsequently categorised according to PseudoCAP (*P. aeruginosa* community annotation project) functions (Figure 3) [43]. Fifteen genes were annotated with at least two functional categories and nine genes were part of regulatory networks (Figure 3), including key global regulators (e.g. *rpoN, mexT*), which might impact multiple processes [44]. Approximately 50% of the non-synonymous mutations occurred in genes that encode proteins associated with virulence (e.g. PilR involved in surface attachment and twitching motility [45, 46]; MigA involved in swarming [47]; ZnuA in zinc homeostasis [48]; PchD for iron acquisition [49]) or antibiotic resistance mechanisms (e.g. OprD: carbapenem resistance [50]; GyrB: fluoroquinolone resistance [51]; FtsI, Mpl: β-lactam resistance [52, 53]; PmrB, ColS: polymyxin resistance [54, 55]; MexA, MexS, MexT: multi-drug efflux pumps [56-59]). The mutations found in genes correlated with antibiotic resistance were also identified in M3L7 isolates collected in 2014 [17] and might help explain the increased resistance described previously in the sub-type compared to other strains [16]. Altogether this analysis suggests that the M3L7 sub-lineage is characterised by mutations in genes that might have aided adaptation to the CF airways prior to dissemination in the CF population [4, 8, 39, 60].

**Figure 3.**
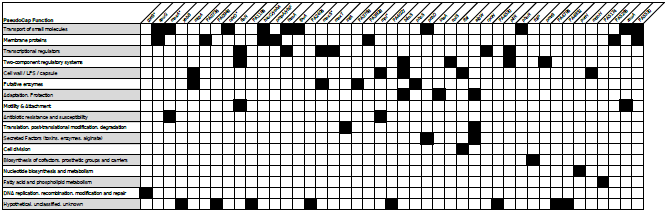
Genes (n=43) with non-synonymous SNPs and indels that characterise the M3L7 sub-lineage. The black squares indicate the PseudoCAP function of the gene. See Additional File 6: Table S2 for detailed information of the specific mutations. *Candidate pathoadaptive genes identified previously [8].

### Genomic variation due to large deletions

The draft genomes that comprise the M3L7 sub-lineage ranged in size from 6.15 to 6.25 Mbp, whereas the M3L1 isolate (AUS970) genome was substantially smaller (6.05 Mbp). All genomes within the M3 clade had an average GC content of 66.5%. Genome reduction is a typical evolutionary process that occurs during adaptation within the CF lungs [61]; therefore, large genomic variations (> 10 kb) were compared between the AUST-02 isolates.

Major differences in genome size were not due to horizontal gene transfer but attributed to several deletion events, summarised in Table 1. For example, one M3L7 isolate (AUS947) lost a large 93 Kbp region encoding a number of virulence genes including *exoY* (a T3SS secreted toxin), a three-part hydrogen cyanide biosynthesis operon, *hcnABC* and part of the *cupA1-A5* chaperone usher fimbrae operon (Table 1). Deletions of varying sizes encompassing *exoY* and *hcnABC* (of the same region) were also missing in three other isolates within the M3 clade (AUS853; AUS854; AUS970) suggesting that there was a selective pressure to lose the functionality of genes encoded at this position.

**Table 1.**
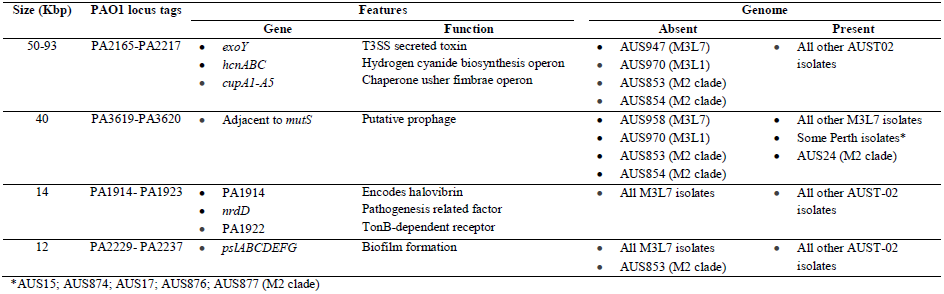
Comparison of deletion events larger than 10 Kb in the AUST-02 genomes.

A 40 Kbp putative prophage was present in five AUST-02 isolates from Perth (AUS15; AUS874; AUS17; AUS876; AUS877) and one isolate from Brisbane (AUS24) of the M2 clade. This prophage is present in nearly all M3L7 genomes (absent in AUS958) and a previous study also observed the presence of the prophage in 9/11 M3L7 isolates collected in 2014 [17]. Therefore, it is most likely that this prophage was present in the last common ancestor of the M3L7 sub-lineage and was vertically inherited. The prophage is inserted immediately adjacent to the *mutS* gene and a search of the putative prophage sequence using the PhaST webtool predicts a complete prophage that is only 9% identical to the *P. aeruginosa* F10 phage [35]. The prophage is flanked by an 18 bp *att* sequence (TCTCTCAGCACACGCC) that delineates the deletion in AUS958.

In conclusion, the persistence of the AUST-02 shared strain in the CF population in Australia has led to the emergence of a monophyletic sub-lineage (M3L7) that is distinct from the M2 sub-lineage of AUST-02. Our WGS analysis demonstrated that M3L7 strains are characterised by mutations in genes that are likely to affect antibiotic resistance and virulence phenotypes. The rapid dissemination of this clinically important sub-type is most likely due to a combination of adaptation to the CF airway microenvironment and transmission between people attending the same CF centre. This work highlights the importance of clinical management in the emergence of shared strains amongst people with CF and provides a framework for future efforts in real-time genomic surveillance to monitor the transmission and pathogenicity of AUST-02 amongst the Australian CF population and to detect newly emergent shared strains.

## Declarations

### Ethics approval and consent to participate

Ethics approval for this project was granted under HREC/07/QRCH/9 and HREC/13/QPCH/127 by The Prince Charles Hospital Human and Research Ethics Committee, Metro North Hospital and Health Service, Brisbane, Queensland, Australia and all participants provided written, informed consent.

### Consent for publication

Not applicable (no individual level patient data contained within this manuscript).

### Availability of data and material

Genome sequence data generated in this study was deposited in the European Nucleotide Archive under study PRJEB14781 with accession identifiers ERS1249679 to ERS1249704. Other AUST-02 isolates are available as part of a separate study (PRJEB21755).

## Author’ s contributions

AST, TJK, DMW, SCB and SAB designed the study. AST, TJK, KAR, DMW and SCB collected, performed genotyping and selected samples for sequencing. BAW and LJS carried out the comparative genome analyses and wrote the original manuscript draft. NLBZ performed BEAST phylodynamic analyses. KRH, NLBZ, IL and SAB contributed to the interpretation of the bioinformatics analyses. KAR and SCB collected information on numbers of CF patients attending TPCH. DMW, SCB and SAB supervised the project. All authors reviewed, edited and approved the final manuscript.

## Acknowledgements

We thank Ms Rebecca Stockwell (Lung Bacteria Group, QIMR Berghofer Medical Research Institute, Brisbane, Australia) for extracting the genomic DNA for WGS.

## Additional Files

**Additional File 1: Table S1.** Details of *Pseudomonas aeruginosa* isolates used in this study.

**Additional File 2: Figure S1.** Phylogeny of sequenced AUST-02 strains showing the M3L7 isolates in relation to other AUST-02 isolates within the major M2 and M3 clades. The Maximum-Likelihood phylogenetic tree was estimated from an alignment of 30,811 core genome SNPs using RAxML. > 70% Bootstrap support (*), 100% bootstrap support (**). Scale indicates branch length representing 10 nucleotide substitutions. M2, *mexZ-2* allele (codon substitution, A38T); M3, *mexZ-3* allele (codon substitution, T12N). Pairs of isolates (collected in 2007 or 2008 and 2011) are indicated by colour. Dotted lines indicate branches that have been shortened and are not to scale. Geographic location of CF treatment centre where each isolate was obtained is shown on the right of the tree.

**Additional File 3: Figure S2.** Growth of the adult CF population at The Prince Charles Hospital between 2001 and 2015.

**Additional File 4: Figure S3.** A resolved phylogeny inferred from 364 core SNPs. The maximum likelihood tree was generated using RAxML with 1000 bootstrap replicates. Bootstrap values > 70% are shown. Scale indicates branch length representing 5 nucleotide substitutions. Non-hypermutator isolates (n=23) sequenced as part of this study were included. Pairs of isolates were collected in 2007 or 2008 and 2011 as indicated by colour. M3, *mexZ-3* allele (codon substitution, T12N); L1, *lasR-1* allele (wild-type); L7, *lasR-7* allele (1 bp deletion, 438delG).

**Additional File 5: Figure S4.** Core SNP based minimum spanning tree depicting the most likely route of transmission. Isolates sequenced in this study included. Two isolates (AUS946 and AUS944) were predicted to be closest to the source. Non-hypermutator isolates sequenced in this study were included. P, patient. Figure generated using Phyloviz [29].

**Additional File 6: Table S2.** Position and details of non-synonymous SNPs (n=35) and indels (n=9) that characterise the M3L7 sub-lineage.

